# Kv2–VAP interactions enhance presynaptic ER and mitochondrial calcium influx and the mobilization of vesicles from the reserve pool

**DOI:** 10.64898/2026.05.04.722784

**Authors:** Cameron D Paton, Michael B Hoppa

## Abstract

In neurons, the endoplasmic reticulum (ER) forms an extensive network that establishes membrane contact sites (MCSs) with various organelles including the plasma membrane (PM). While MCSs are known to regulate lipid exchange and Ca^2+^ signaling, their specific roles in synaptic transmission remain poorly understood. Here, we demonstrate that the ER resident proteins VAPA and VAPB are essential for organizing presynaptic Ca^2+^ exchange and mobilizing synaptic vesicles. We show that the loss of VAP impairs Ca^2+^ loading into both the ER and mitochondria during electrical activity. This regulation occurs primarily through VAP interactions with voltage-gated potassium channels (Kv2) at the PM. Our data suggest that the Kv2-VAP complex organizes presynaptic Ca^2+^ signaling outside of the active zone. Without this scaffold, synaptic vesicles become trapped in the reserve pool and fail to participate in exocytosis. These findings reveal a novel role for Kv2-VAP MCSs in coordinating organelle Ca^2+^ signaling and the synaptic vesicle cycle.

## Introduction

Advanced 3D ultrastructural visualization has prompted a new conceptual framework for the structural organization of the endoplasmic reticulum (ER) in neurons. As the largest membrane-bound organelle in eukaryotic cells, the ER exhibits a complex morphology defined by two predominant motifs: dense interlocking tubules (10-100nm diameter) and flattened sheets or cisternae (1–5). This structural distinction is highly polarized within neurons. In the soma and dendrites, the ER is primarily organized into small cisternae that form ladder-like structures, which are interspersed with periodically arranged membrane contact sites (MCSs) that coordinate Ca^2+^ signaling (6). In contrast, the axonal ER consists of smooth narrow tubules that extend into terminal processes. At presynaptic boutons, these tubules transition into a highly branched network that physically interacts with mitochondria and synaptic vesicles (4, 5). While the physical architecture of the axonal ER is well-documented, the functional mechanisms by which these inter-organelle interactions support synaptic transmission remain an active area of inquiry.

A primary function of the smooth ER is the dynamic regulation of Ca^2+^. The ER serves as the largest intracellular Ca^2+^ reservoir, with a lumenal concentration of approximately 150μM, over three orders of magnitude higher than 100nM in the cytosol (7). Despite its limited volume within the axon, the ER can act as both a sink and source to modulate local Ca^2+^ transients. During neuronal activity, presynaptic voltage-gated Ca^2+^ channels (Cavs) in nerve terminals open, elevating cytosolic Ca^2+^ within microdomains (8, 9). The ER rapidly sequesters Ca^2+^ through the activity of sarco-endoplasmic reticulum Ca^2+^ adenosine ATPases (SERCAs), working in tandem with membrane Ca^2+^ ATPases (PMCAs) to restore homeostasis (10). Conversely, the ER can amplify signals, by releasing Ca^2+^ through ryanodine receptors (RyRs) or IP_3_ receptors (IP_3_Rs) (10–13). Although evidence suggests the ER is functionally-coupled to Cavs at the plasma membrane (14–15), the precise molecular bridge linking ER Ca^2+^ handling to the machinery of synaptic transmission is not yet understood.

The functional role of the ER in the synapse likely depends on its physical proximity to other organelles at MCSs, where membranes are held within <30nm of each other (16, 17). A key candidate for mediating these interactions is the ER-resident protein VAP (Vesicle Associated Membrane Protein-Associated Protein). Highly conserved and ubiquitously expressed, VAP was originally considered as a potential partner of the SNARE complex, where its disruption was shown to impair synaptic transmission (18). In vertebrates, two isoforms, VAPA and VAPB (19), that have been shown to bind to many intracellular protein targets containing a highly conserved ‘two phenylalanine in an acidic tract’ (FFAT) motif (20–22). The clinical importance of the pathway is underscored by VAPB mutations linked to Amyotrophic Lateral Sclerosis (ALS), which present with clear synaptic deficits (23, 24)

VAP isoforms primarily localize to two critical presynaptic hubs: ER-mitochondria contacts (via PTPIP51) and ER-PM contacts (via voltage-gated Kv2 K^+^ channels) (25). We previously established that Kv2.1 channels are vital for synaptic function, as their loss impairs Ca^2+^ influx into the axon and reduces exocytosis by ∼50% (26). However, whether VAP serves as the essential tether that couples Kv2-mediated Ca^2+^ dynamics to transmission and the synaptic vesicle cycle remains to be determined.

In this study, we investigate how VAP-mediated MCSs coordinate Ca^2+^ handling and vesicle mobilization in the presynaptic bouton. We show that VAPA and VAPB cluster Kv2.1 channels in hippocampal neurons and that the loss of VAP severely impairs synaptic transmission. Specifically, VAP deficiency decouples both the ER and mitochondria from cytosolic Ca^2+^ influx during electrical activity—a phenotype mirrored by Kv2 depletion but not by loss of the mitochondrial tether PTPIP51. Finally, we demonstrate that VAP is required to mobilize synaptic vesicles from the quiescent reserve pool. Together, our results identify VAP-mediated ER-PM contact sites as a master regulator of presynaptic Ca^2+^ exchange and vesicle availability.

## Results

### Kv2.1 channel clustering is regulated by VAP expression

Kv2 channels are highly expressed throughout the brain (27), and exist in diffuse or clustered populations, where they have different roles. Diffuse Kv2 play a conducting role in membrane repolarization in the soma (27–29), while clustered Kv2 are non-conductive (30). In heterologous cells, VAP proteins colocalize at MCSs with Kv2 and are necessary for clustering (31, 32). Whether VAP isoforms regulate Kv2 clustering in neurons has never been tested. Neurons also express other ER proteins with FFAT-binding motifs, such as MOSPD-2 (33–35). This raises an important question: Does Kv2 clustering require VAP isoforms specifically, or can other FFAT proteins mediate this process? To visualize Kv2.1 clusters, we utilized a fluorescent chimera (mGreenLantern-Kv2.1). When expressed in neurons, mGreenLantern-Kv2.1 forms a pattern of clusters, each up to a micron in size, across the somatic compartment as well as the axon initial segment (Fig 1A). To test the molecular function of VAP isoforms in the axon and presynaptic boutons, we developed a bicistronic CRISPR sgRNA construct to deplete both isoforms of VAP (VAPA/B), with quantification shown in Supplementary Figure S1 (36). Loss of VAPA/B leads to a redistribution of Kv2.1 and loss of clustering (Fig 1B). To determine the degree of clustering, we took sa ingle z-plane of the soma and quantified the coefficient of variation (CV) along a line scan around the soma (Fig 1C-D, F). Overall, clustering was impaired in VAPA/B KD neurons, compared to control neurons. We were able to restore Kv2.1 clustering in knockdown neurons by expressing a human isoform of VAPB, hVAPB (Fig 1F). Interestingly, we found that Kv2.1 clustering remained visible in the axon initial segment (AIS) in VAPA/B knockdown neurons (Fig 1B, arrowhead), consistent with prior reports of FFAT independent localization at the AIS (37) and possibly the result of slower protein turnover (38, 39). We only quantified clustering in the soma at ER-PM MCSs, as the signal in axons is very dim due to the lower number of proteins, likely due to the ∼30 nm diameter of axonal MCSs (5). We hypothesize that this organization extends into the bouton, with accumulating evidence showing enrichment of Kv2 channels in distal the axon and boutons (26, 40, 41). Taken together, these results suggest that VAP is necessary for Kv2.1 clustering in neurons.

**Figure 1.**
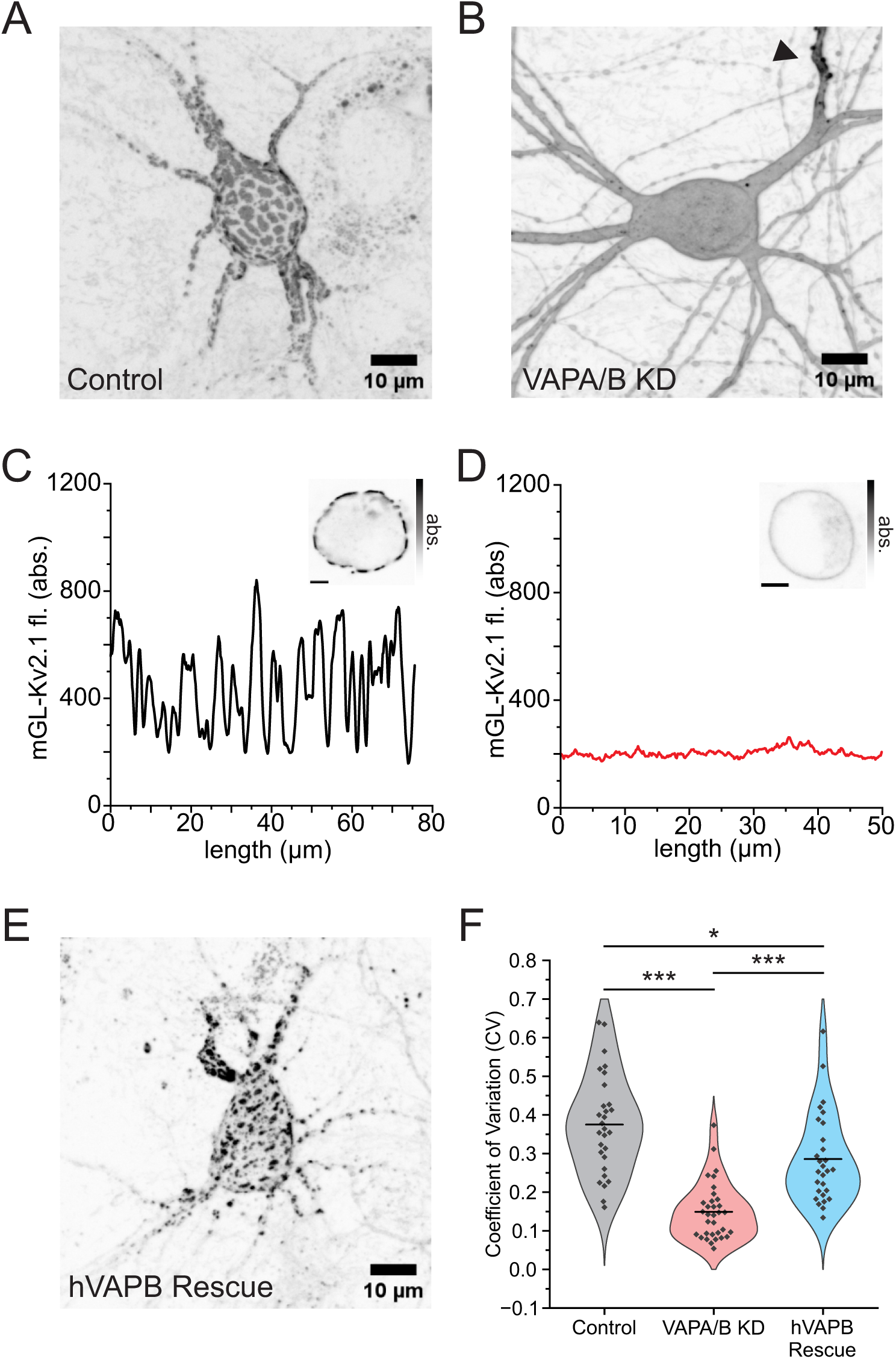
VAPA/B organize Kv2.1 clustering in the soma. (A) Representative image of mGreenLantern-Kv2.1 under the human synapsin 1 promoter in a hippocampal neuron. (B) After VAPA/B knockdown (KD) using a CRISPR sgRNA against both VAP isoforms, Kv2.1 clustering is lost in the soma but remains in the axon initial segment (arrowhead), and sparse localization is present in distal processes. (*C* and *D*) To determine whether VAP isoforms contribute to Kv2.1 clustering, we used the coefficient of variation from a line scan taken from single z plane through the soma. Inset, single plane used to calculate coefficient of variation (CV). Scale bar, 5µM. (E) Representative image of mGreenLantern-Kv2.1 in a VAPA/B KD hippocampal neuron coexpressing human VAPB (hVAPB). (F) Expression of human VAPB (hVAPB) in the KD background is able to partially rescue Kv2.1 clustering above KD levels, but not to control levels (control neurons, CV, 37.5 ± 2 %, n = 29; VAPA/B KD neurons, CV, 14.9 ± 1 %, n = 33; hVAPB neurons, CV, 28.6 ± 2 %, n = 28;\*\**P*<0.01, \*\*\**P*<0.001, One-way ANOVA, *post hoc* Tukey’s test).

### Loss of ER Proteins VAPA/B impairs synaptic transmission

To test the role of VAP in synaptic transmission, we measured exocytosis following stimulation using vGlut-pHluorin (vG-pH) quantified as a percentage of the total vesicle pool identified using NH_4_Cl as previously described (42, 43). We found a 51% impairment in exocytosis in VAPA/B KD neurons compared to control neurons (control, 10.5 ± 1 %; VAPA/B KD neurons, 3.8 ± 1 %; *P* < 0.001; all statistics are mean ± SEM unless specified otherwise) (Fig 2A-B). These findings mimic the exocytosis impairment in Kv2.1 KD neurons (26), suggesting that the formation of Kv2.1/VAP MCS between the PM and ER are required for synaptic function. To test whether VAP proteins have isoform-specific roles in exocytosis, we expressed sgRNA-resistant human isoforms of VAPA (hVAPA) or VAPB (hVAPB) (Fig 2A-B, C-D, and respectively) in KD neurons. Both hVAPA and hVAPB alone were able to rescue exocytosis to control levels (Fig 2B, D). VAP proteins bind their partners through an amino acid motif location in the N-terminal major sperm protein domain (21). Kv2 channels form non-conducting clusters via their C-terminus, which contains a non-canonical FFAT motif (two phenylalanines in an acidic tract) that binds VAP proteins (29, 31, 32). To test whether VAP’s role in synaptic function depends on this FFAT-mediated binding, we generated a VAPB mutant (K87D/M89D) that disrupts FFAT binding interactions (44). We were unable to rescue exocytosis when we expressed VAPB K87D/M89D in KD neurons (control neurons, 9.0 ± 1 % exocytosis; hVAPB K87D/M89D neurons, 5.4 ± 1 % exocytosis; *P* < 0.05) (Fig 2F). These results demonstrate that VAP proteins play an essential role in supporting synaptic transmission, that the two isoforms are redundant, and that VAP requires FFAT binding to support synaptic function.

**Figure 2.**
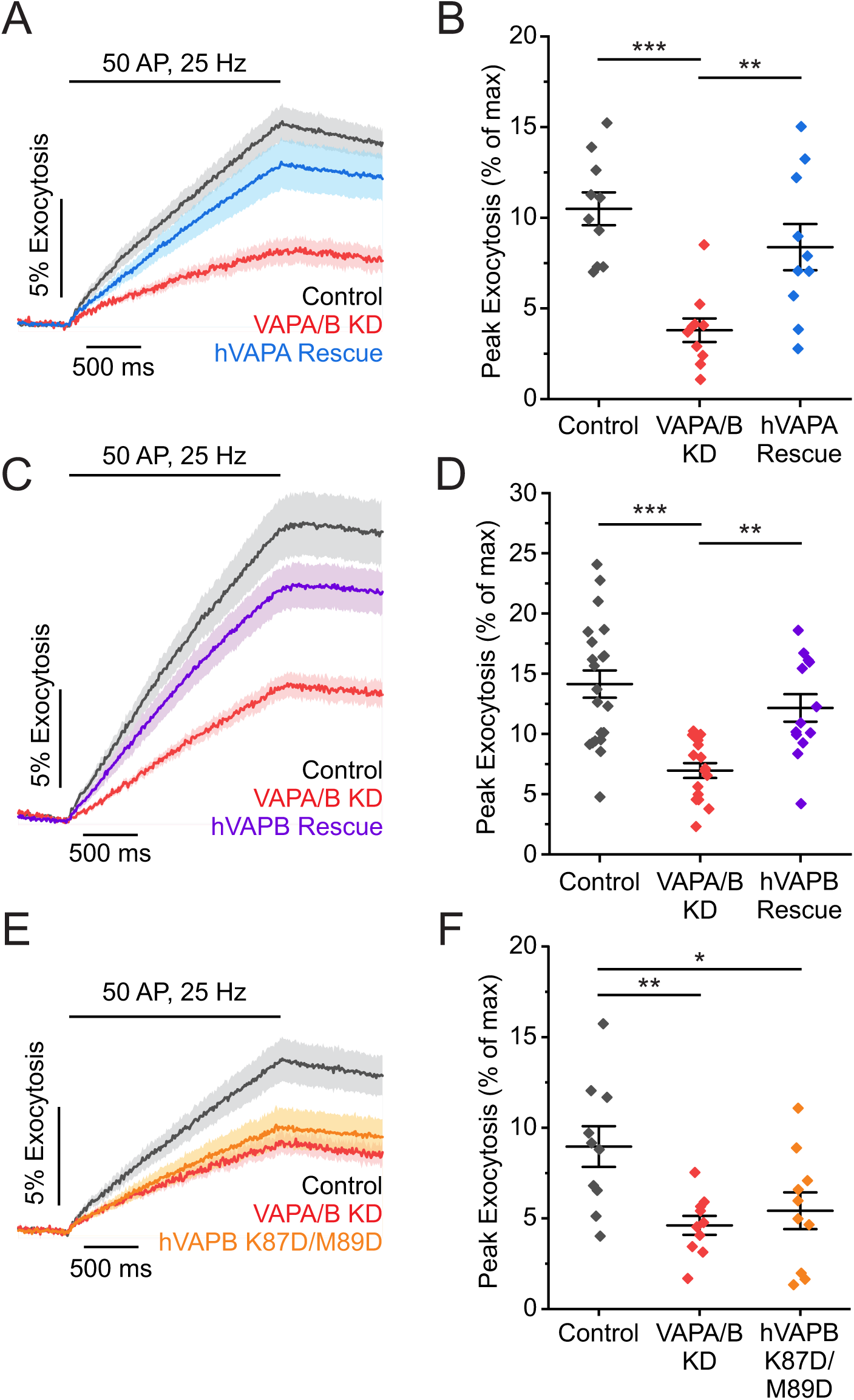
Loss of VAPA/B impairs vesicle fusion and requires key residues that coordinate FFAT binding to partners like Kv2.1. (*A* and *B*) Average fluorescent traces of vGlut-pHluorin (vG-pH) (*A*) and quantification of peak fluorescence (*B*) for control, VAPA/B KD, and hVAPA rescue neurons. hVAPA rescues exocytosis impairment in KD background (control, n = 10; VAPA/B KD, n = 10; hVAPA rescue, n = 10; \*\**P*<0.01, \*\*\**P*<0.001, One-way ANOVA *post hoc* Tukey’s test). (*C* and *D*) Average fluorescent traces of vG-pH (*C*) and quantification of peak fluorescence (*D*) for control, VAPA/B KD, and hVAPB rescue neurons. hVAPB rescues exocytosis impairment in the KD background (control neurons, n =21; VAPA/B KD neurons, n = 16; hVAPB rescue neurons, n =13; \*\**P*<0.01, \*\*\**P*<0.001, One-way ANOVA *post hoc* Tukey’s test). (*E* and *F*) Average fluorescent traces of vG-pH (*E*) and quantification of peak fluorescence (*F*) for control, VAPA/B KD, and hVAPB K87D/M89D rescue neurons (control, n = 10; VAPA/B KD, n = 10; hVAPB K87D/M89D rescue, n = 10; \**P*<0.05, \*\**P*<0.01, One-way ANOVA *post hoc* Tukey’s test).

### VAPA/B KD impairs ER lumen Ca^2+^ filling during electrical activity

Previous studies revealed a non-conducting role for Kv2.1 in regulating ER Ca^2+^ filling during electrical activity or following ER store depletion (26). Given the physical association between these two proteins, we hypothesize that VAP facilitates ER Ca^2+^ uptake through its interaction with Kv2. To test this, we examined whether depleting VAPA/B disrupts Ca^2+^ dynamics in both the cytosol and the ER lumen. We measured cytosolic Ca^2+^ using the genetically encoded Ca^2+^ indicator, jGCaMP8f (Fig 3A)(45). Loss of VAP resulted in no significant change to action potential (AP)-evoked cytosolic Ca^2+^ influx (P = 0.96), suggesting that the overall balance of cytosolic Ca^2+^ entry and clearance remains intact (Fig3B). We also found no change in peak AP-evoked cytosolic Ca^2+^ influx (*P* = 0.96) (Fig 3C). We then monitored ER Ca^2+^ influx using the genetically encoded ER lumen Ca^2+^ indicator, ER-GCaMP-150 as seen in Figure 3D (7). In contrast to the cytosolic results, we found a 56% impairment in ER Ca^2+^ uptake in VAPA/B KD neurons (control, 21.5 ± 4 % ΔF/F; VAPA/B KD, 9.4 ± 2 % ΔF/F; *P* < 0.05) (Fig 3E-F). These findings largely parallel the ER Ca^2+^ influx impairments seen when Kv2 is depleted (26), although a notable difference exists: during train stimulation, loss of Kv2.1 – but not VAP – also leads to a decrease in cytosolic Ca^2+^ influx (Fig S2). This could point to a unique role for Kv2 channels to recruit Cavs (14, 46) and RyR (47, 48). Overall, our results indicate VAP-mediated ER-PM MCSs create a uniquely organized site of Ca^2+^ entry that is preferentially coupled to organelle sequestration rather than the primary synaptic vesicle release machinery.

**Figure 3.**
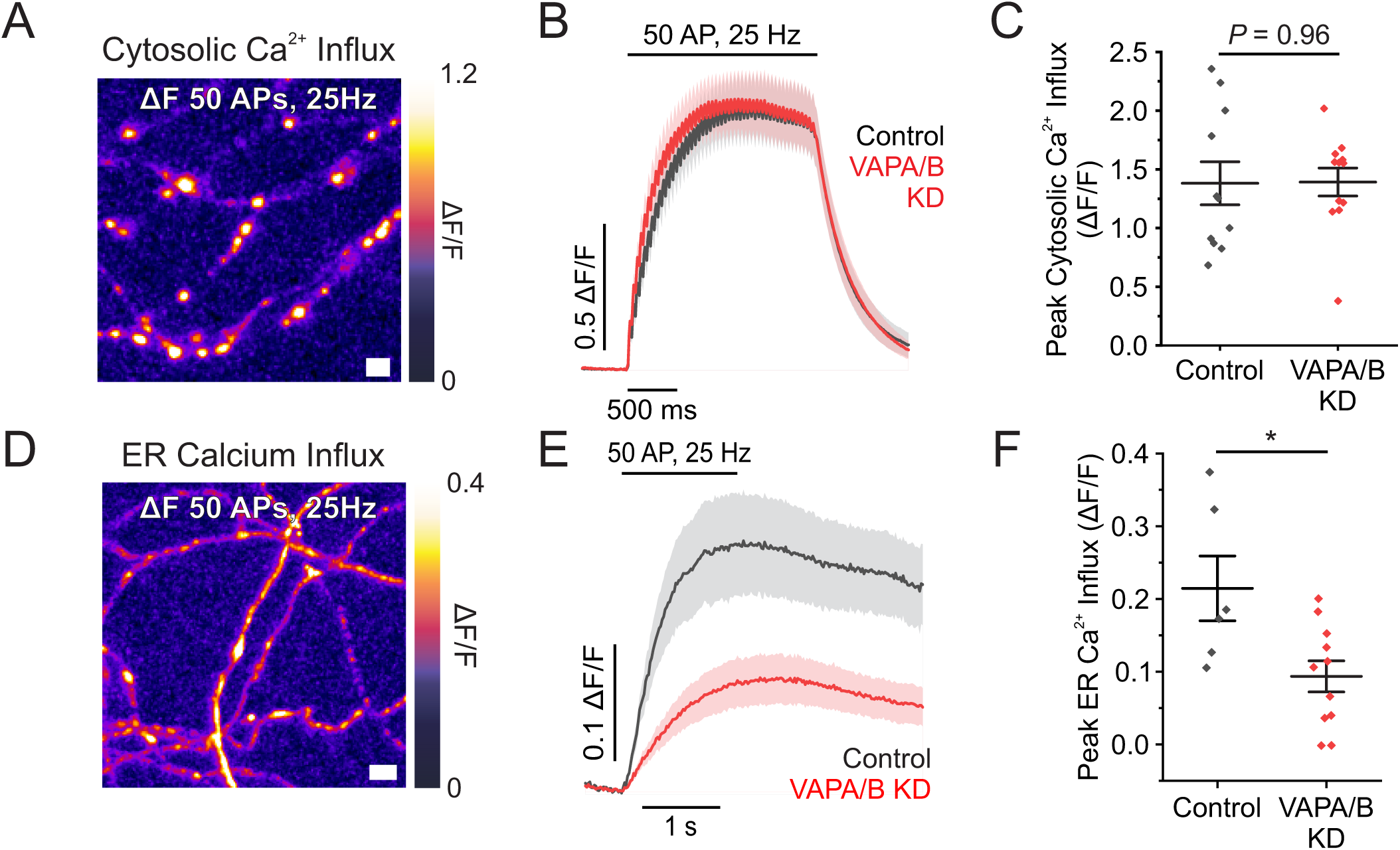
Loss of VAP decouples activity dependent ER Ca^2+^ uptake from cytosolic Ca^2+^ influx. (A) Representative image of the change in fluorescence of jGCaMP8f in response to a train of stimulation. Scale bar 5 µm. (B) Average fluorescent traces of jGCaMP8f (*B*) and quantification of peak fluorescence (*C*) for control and VAPA/B KD neurons (control, n = 11; VAPA/B KD, n = 12; *P* = 0.96, Student’s *t-*test). (D) Representative image of the change in fluorescence of ER-GCaMP6-150 in response to a train of stimulation. Scale bar 5 µm. (E) Average fluorescent traces of ER-GCaMP6-150 (*E*) and quantification of peak fluorescence (*F*) for control neurons and VAPA/B KD neurons (control, n = 6; VAPA/B KD, n = 11; \**P*<0.05, Student’s *t-*test).

### VAPA/B supports Ca^2+^ influx to the mitochondria, which is dependent on Kv2 mediated MCS

We next investigated whether VAP-mediated organization extends to other organelles, specifically mitochondria. During electrical activity, mitochondria rapidly sequester Ca^2+^, which can be utilized for citric acid dehydrogenase enzyme function to generate ATP (49). This has been shown to help meet high presynaptic demand for ATP in supporting the synaptic vesicle cycle (50). To determine if the loss of VAP alters mitochondria Ca^2+^ signaling, we used a mitochondrial matrix-localized Ca^2+^ indicator as previously described, mito^4x^-GCaMP6f (50). In VAPA/B KD neurons, we found an 87% decrease in mitochondrial Ca^2+^ influx (control, 67.2 ± 7% ΔF/F; VAPA/B KD, 8.6 ± 3% ΔF/F; *P* < 0.001) (Fig 4A-B). Presynaptically VAPA/B can form MCSs with mitochondria through PTPIP51 (25) and with the PM through Kv2.1 (31, 32). To determine which interaction is critical for mitochondrial Ca^2+^ filling, we performed individual knockdowns of these partners. Interestingly, knockdown of PTPIP51 did not impair mitochondrial Ca^2+^ influx (*P* = 0.12) (Fig 4D. In contrast, Kv2.1 knockdown impaired mitochondrial Ca^2+^ influx by 58% (control, 42.8 ± 4 % ΔF/F; Kv2.1 KD, 18.0 ± 3 % ΔF/F; *P* < 0.001) (Fig 4F). Together, these data suggest that VAP-Kv2 MCSs, rather VAP-PTPIP51 MCSs, are the major regulators of Ca^2+^ influx into presynaptic mitochondria.

**Figure 4.**
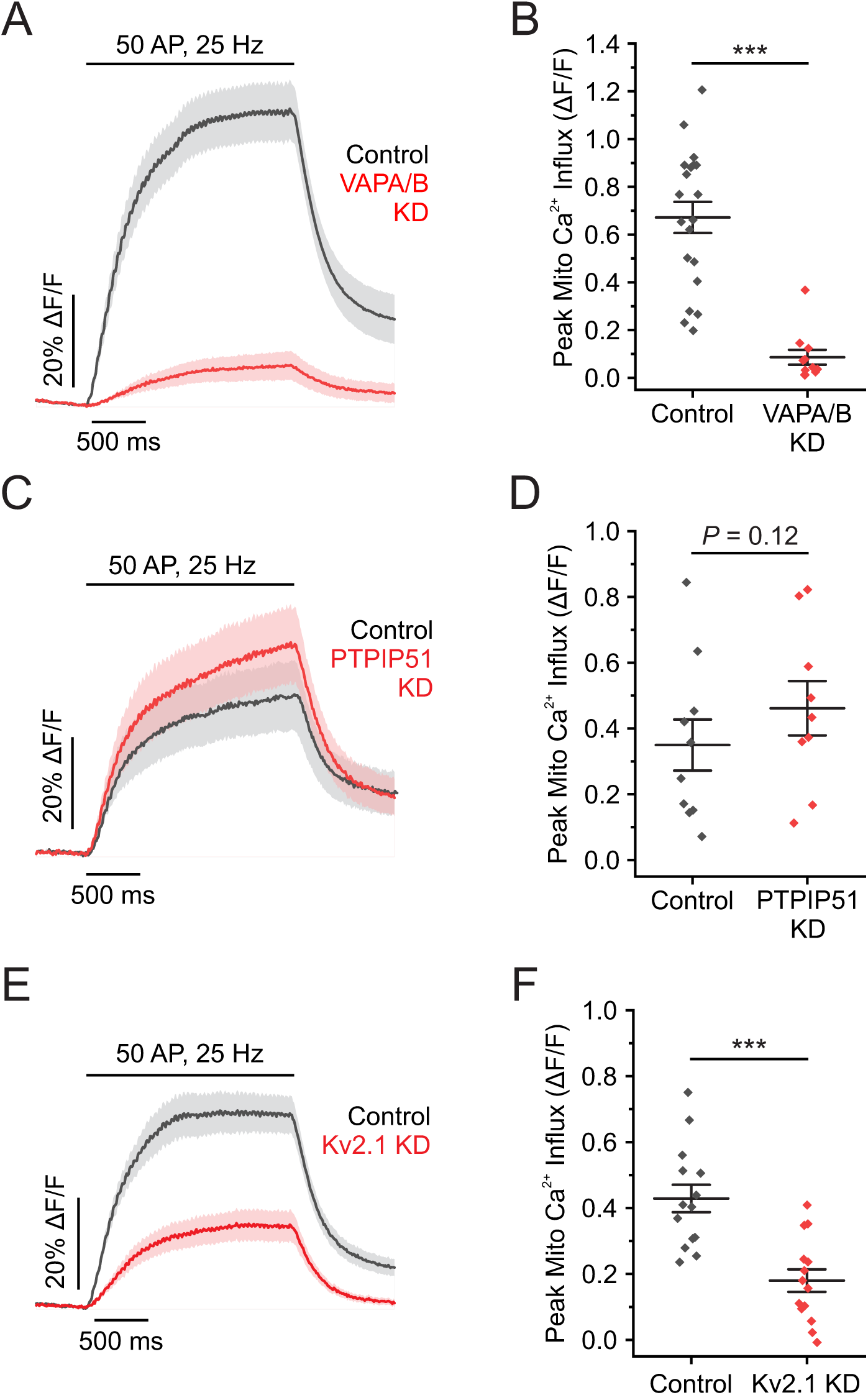
VAPA/B MCS with Kv2 support mitochondrial Ca^2+^ influx. (*A* and *B*) Average fluorescent traces of mito^4x^-GCaMP6f (*A*) and quantification of peak fluorescence (*B*) for control and VAPA/B KD neurons (control, n = 20; VAPA/B KD, n = 11; \*\*\**P*<0.001, Student’s *t-*test). (*C* and *D*) Average fluorescent traces of mito^4x^-GCaMP6f (*C*) and quantification of peak fluorescence (*D*) for control and PTPIP51 KD neurons (control, n = 10; PTPIP51 KD, n = 9; *P*=0.12, Student’s *t-*test). (*E* and *F*) Average fluorescent traces of mito^4x^-GCaMP6f (*E*) and quantification of peak fluorescence (*F*) for control and Kv2.1 KD neurons (control, n = 14; Kv2.1 KD, n = 14; \*\*\**P*<0.001, Student’s *t-*test).

### Loss of VAPA/B impairs mitochondrial access to Ca^2+^ from Cavs

To sequester Ca^2+^ from the cytosol, mitochondria in the axon rely on the subunit MICU3, which adapts the mitochondria Ca^2+^ uniporter (MCU) to facilitate Ca^2+^ uptake from the cytosol (49–51). This suggests that the impairment found in both organelles with the loss of VAP-Kv2 MCSs (ER and mitochondria) are simply correlated and not causative. To confirm that mitochondria are not receiving Ca^2+^ from the ER, we applied 50 µM cyclopiazonic acid (CPA) to block ER Ca^2+^ signaling by inhibiting SERCA pumps and depleting ER luminal Ca^2+^(26). CPA treatment had no effect on mitochondria Ca^2+^ influx (*P* = 0.87) (Fig 5B). Additionally, we applied 10 µM dantrolene, a selective RyR1,3 antagonist to block potential sources of Ca^2+^-induced Ca^2+^ release. Dantrolene treatment caused a small decrease in mitochondrial Ca^2+^ influx that became more pronounced the longer stimulation continued (Fig 5D). Together, our results indicate that mitochondria Ca^2+^ influx during stimulation is facilitated primarily by VAP-Kv2 MCSs and potentially by RyRs during repeated stimulation, but not from VAP-PTPIP51 MCSs or dependent on the ER as a primary source of Ca^2+^.

**Figure 5.**
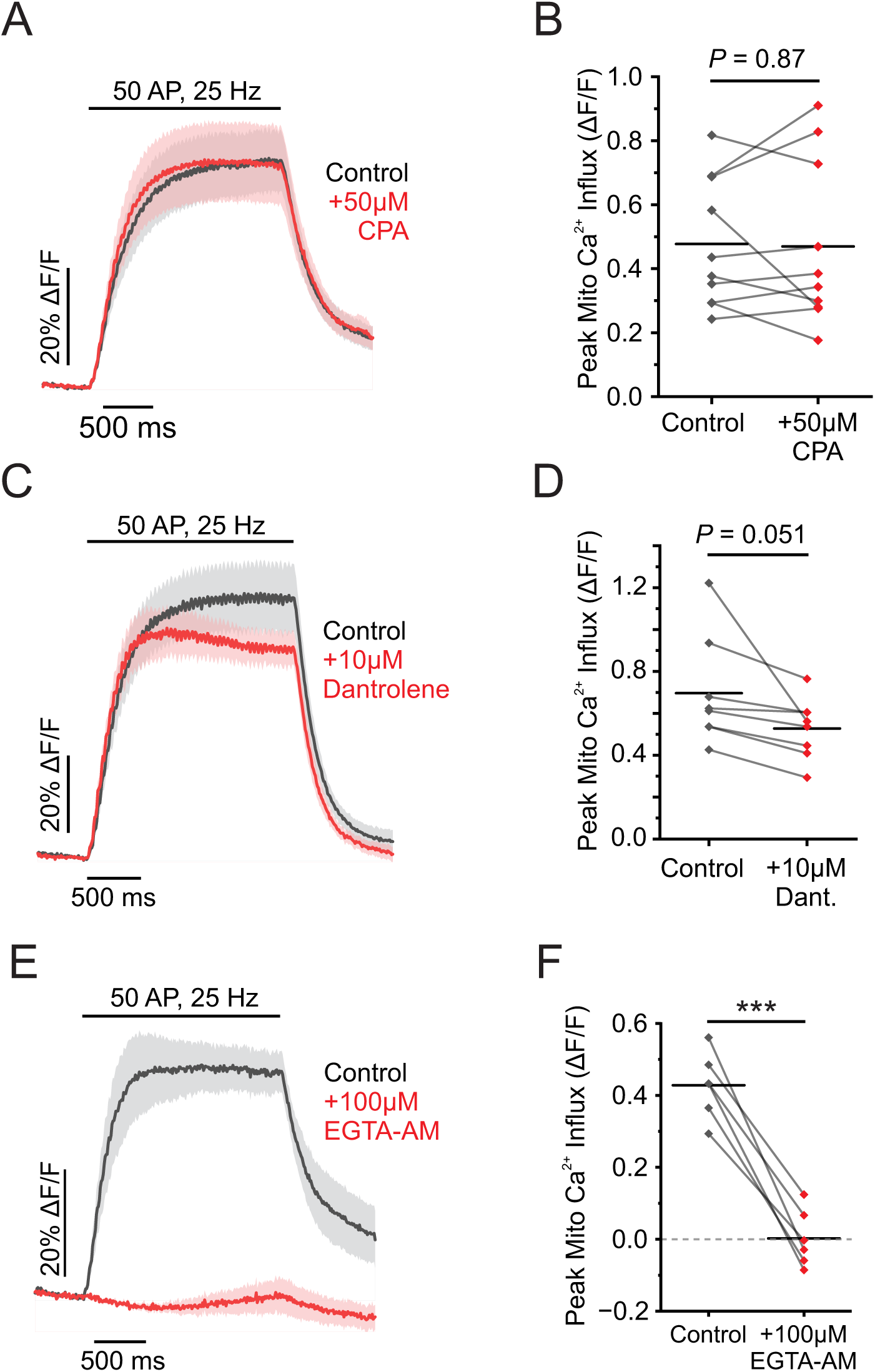
Mitochondria are loosely coupled to ER Ca^2+^ release through Ryanodine receptors. (*A* and *B*) Average fluorescent traces of mito^4x^-GCaMP6f (*A*) and quantification of peak fluorescence (*B*) for neurons ± 50µM CPA (n = 10; *P* = 0.87, paired *t-*test). (*C* and *D*) Average fluorescent traces of mito^4x^-GCaMP6f (*C*) and quantification of peak fluorescence (*D*) for neurons ± 10µM dantrolene (n = 8; *P* = 0.051, paired *t-*test). (*E* and *F*) Average fluorescent traces of mito^4x^-GCaMP6f (*E*) and quantification of peak fluorescence (*F*) for neurons ± 100µM EGTA-AM (n = 6, \*\*\**P* < 0.001, paired *t-*test).

Kv2.1 channels were previously shown to colocalize with L-type voltage gated Ca^2+^ channels (Cav) in the soma and dendrites. Furthermore, this work identified a functional interaction between Kv2.1, L-type Ca^2+^ channels (Cav1.2 and Cav1.3), and RyRs in the generation of somatodendritic Ca^2+^ signals (14). The organization of these proteins we believe extends beyond the soma of neurons and into boutons where they contribute to vesicle fusion machinery. Recent work from *C. elegans* supports this idea by demonstrating that L-type voltage gated Ca^2+^ channels alone are not sufficient for vesicle fusion, but instead rely on functional coupling to RyR to fuse synaptic vesicles (13). These findings suggest that Kv2.1 channels may be capable of organizing sites of Ca^2+^ influx that supports organelle Ca^2+^ uptake and contribute to vesicle fusion. This perhaps explains why the effect of dantrolene seems to peak at the end of train stimulation, after elevations in cytosolic Ca^2+^ stimulate Ca^2+^ release from the ER to the mitochondria. To determine how tightly-coupled mitochondria are to sources of Ca^2+^, we tested their sensitivity to EGTA treatment. If they are tightly-coupled, we expect them to be EGTA-insensitive, similar to functional coupling between Cav2 channels and synaptic vesicles in the active zone (52, 53). Treatment with 100 µM EGTA-AM, a cell-permeable Ca^2+^ chelator, strongly blocked mitochondrial Ca^2+^ influx (99%), supporting loose coupling between these Ca^2+^ sources and mitochondria (control neurons, 42.8 ± 4 % ΔF/F; EGTA-treated neurons, 0.2 ± 3 % ΔF/F; *P* < 0.001) (Fig 5F). Together, this data suggests two conclusions: first, mitochondria receive Ca^2+^ primarily by diffusion from the cytosol rather than through physical coupling; and second, this cytosolic Ca^2+^ source is spatially organized by VAP through its binding with Kv2.

### Loss of VAPA/B does not affect mitochondria localization in presynaptic terminals

While VAP seems to only affect Ca^2+^ handling through binding interactions with Kv2, it is also possible that VAP itself localizes mitochondria to presynaptic terminals. A recent study identified that VAP isoforms play a critical role in mitochondrial localization in dendrites but not in presynaptic compartments (36). The ER and mitochondria are tightly associated with each other in most cells studied, and form MCSs in mammalian cells (5, 25). To determine whether loss of VAP affects the positioning and distribution of mitochondria in presynaptic boutons, we expressed fluorescent reporters for mitochondria, mitoGFP, and for presynaptic boutons, mRuby-Synapsin, and quantified the fluorescence intensity along segments of axon (Fig 6A) in control and VAPA/B KD neurons. We found a slightly higher correlation of mitochondria at KD synapses (control neurons, Spearman’s ρ = 0.48 ± 0.02; VAPA/B KD neurons, Spearman’s ρ = 0.58 ± 0.02; *P* < 0.001) (Fig 6B). To ensure that the higher correlation was not due to changes in the distribution of mitochondria or synapses along axonal segments, we measured their respective distributions and found that there was no difference in the distribution of either (*P* = 0.12 and *P* = 0.47, respectively) (Fig 6C-D). Mitochondria size was also unchanged, by comparing the full width at half-max (*P* = 0.48) (Fig 6E). Finally, we investigated whether the loss of VAP proteins affects mitochondrial proximity to synapses. We found that there was no change to the percentage of synapses containing proximal mitochondria (±1µm) between control and VAPA/B knockdown neurons using an overlap measurement (see methods; *P* = 0.67) (Fig 6F), which is in close agreement with prior measurements (54). Taken together, we determined that the loss of VAP isoforms does not perturb mitochondria localization presynaptically.

**Figure 6.**
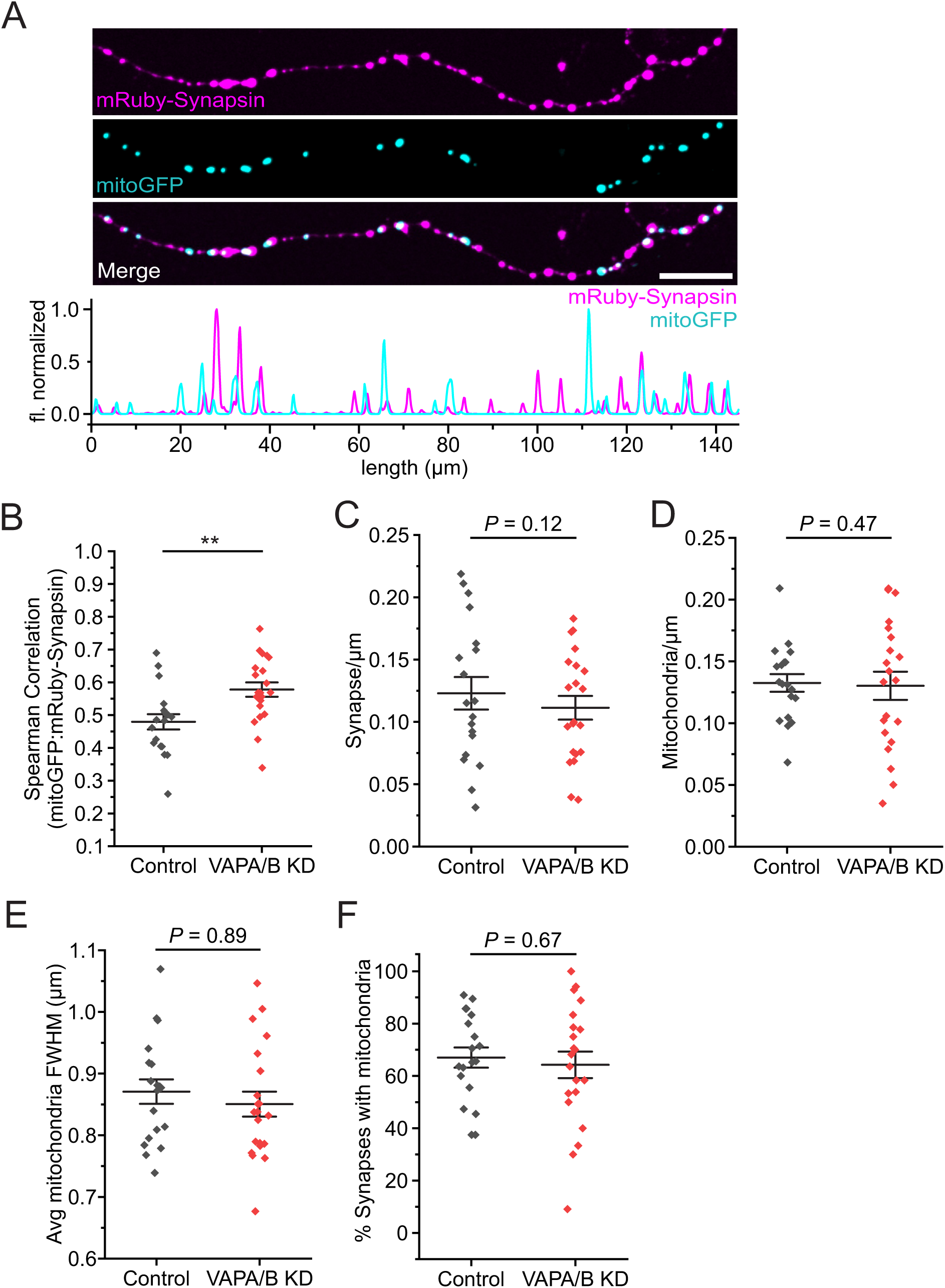
Loss of VAP does not affect mitochondria distribution at synapses. (A) Representative projection (∼2um in volume) line scan along an isolated axon showing mRuby-synapsin fluorescence (top), mitoGFP fluorescence (middle), and merged image with normalized scan (bottom). Scale bar 10µm. (B) Spearman correlation for mRuby-Synapsin and mitoGFP fluorescence in control and VAPA/B KD neurons (control, n = 19; VAPA/B KD, n = 21; Student’s *t-*test, \*\**P* < 0.01). (C) Synapse distribution marked by mRuby-synapsin fluorescence along stretches of axons (control, n = 19; VAPA/B KD, n = 21; *P* = 0.47, Student’s *t-*test). (D) Mitochondria distribution marked by mitoGFP fluorescence along stretches of axon (control, n = 19; VAPA/B KD, n = 21; *P* = 0.12, Student’s *t-*test) (E) Average mitochondria full-width at half-max (FWHM) (control, n = 19; VAPA/B KD, n = 21; *P* = 0.89, Student’s *t-*test) (F) Percent of synapse with mitochondria within 1µm of peak mRuby-Synapsin signal. (control, n = 19; VAPA/B KD, n = 21; *P* = 0.67, Student’s *t-*test).

### VAPA/B does not affect activity-dependent ATP production

In dendrites, VAP proteins contribute to mitochondria Ca^2+^ influx during long durations of synaptic activity to support plasticity-induced changes to dendritic spine morphology and maintenance through ATP generation (36). Given the impairments that we have observed in mitochondrial Ca^2+^ influx, we hypothesized that decreased mitochondrial Ca^2+^ influx may also impair mitochondria’s ability to generate on-demand ATP to support vesicle cycling. One of the most sensitive measurements to changes in ATP levels in synaptic physiology is the rate of vesicle retrieval during endocytosis (55). To test whether there are impairments in on-demand ATP production in VAPA/B KD neurons, we measured endocytosis using vG-pH following a prolonged stimulation, 600AP at 10Hz as previously described (56). After stimulation, the vG-pH signal decays following a single exponential which allows us to compare time constants, τ (57), as a proxy for endocytic rates. To isolate the contribution of mitochondria to on-demand ATP production, we used a modified Tyrode’s imaging solution in which glucose is substituted for lactate and pyruvate (50). We found that endocytic rates (τ) are similar in lactate and pyruvate in control and VAPA/B KD (*P* = 0.28) (Fig S2C), and can be blocked by addition of Oligomycin to show that it’s specific to cellular respiration. This data suggests that while VAP MCSs contribute to mitochondria Ca^2+^ influx, they don’t seem to play a role in the ability for axonal mitochondria to generate activity-dependent ATP.

### Loss of VAPA/B impairs the proportion of vesicles available for release during high frequency stimulation

Synapses contain three distinct vesicle pools —readily releasable, recycling, and reserve. Synaptic strength during sustained activity depends on mobilizing vesicles from a robust recycling pool to replace vesicles that fuse from the readily releasable pool. While prior work has focused on vesicle-associated Ca²⁺ sensors and molecular distinctions between pools (58–60), we hypothesized that additional Ca²⁺ sources, including internal ER stores, contribute to vesicle mobilization. Because we found that organelle Ca²⁺ signaling is disrupted in presynaptic boutons after VAPA/B depletion, we asked whether VAP-dependent membrane contact sites (MCSs) help maintain access to fusion-competent vesicle pools.

To test whether VAP-dependent membrane contact sites (MCSs) contribute to synaptic vesicle mobilization, we quantified vesicle pool fractions using vG-pH during extensive stimulation (>100 s at 10 Hz) in the presence of bafilomycin A1, an inhibitor of the vesicular V-type ATPase (V-ATPase). Under control conditions, following exocytosis synaptic vesicle components are retrieved by endocytosis and the vesicle lumen is reacidified by the V-ATPase; thus, during stimulus trains, vG-pH fluorescence reflects the net balance of exocytosis, endocytosis, and reacidification. In contrast, when reacidification is blocked by bafilomycin, internalized vesicles remain alkaline and fluorescent, such that the vG-pH signal reports cumulative exocytosis. With extensive stimulation in bafilomycin (>100 s), all fusion-competent vesicles undergo exocytosis at least once and remain unavfluorescent after retrieval, producing a plateau that reports the total recycling pool size (Fig. 7A). To quantify the reserve pool, we then applied 50 mM NH₄Cl to alkalinize vesicles that had not fused, revealing the remaining, non-recycled population as previously described (61, 62). Comparing the bafilomycin plateau to the NH₄Cl-evoked total signal allowed us to determine the fraction of vesicles in the recycling versus reserve pools (Fig. 7B). Using this approach, we found that the recycling pool fraction in VAPA/B knockdown neurons was 36% smaller than in controls (control: 64.7 ± 2% of total vesicles; VAPA/B KD: 41.3 ± 4% of total vesicles; P < 0.001) (Fig. 7C-D). Together, these results indicate that VAPA/B depletion impairs vesicle mobilization into the recycling pool, consistent with the substantial reduction in vesicle fusion observed in VAPA/B KD boutons.

**Figure 7.**
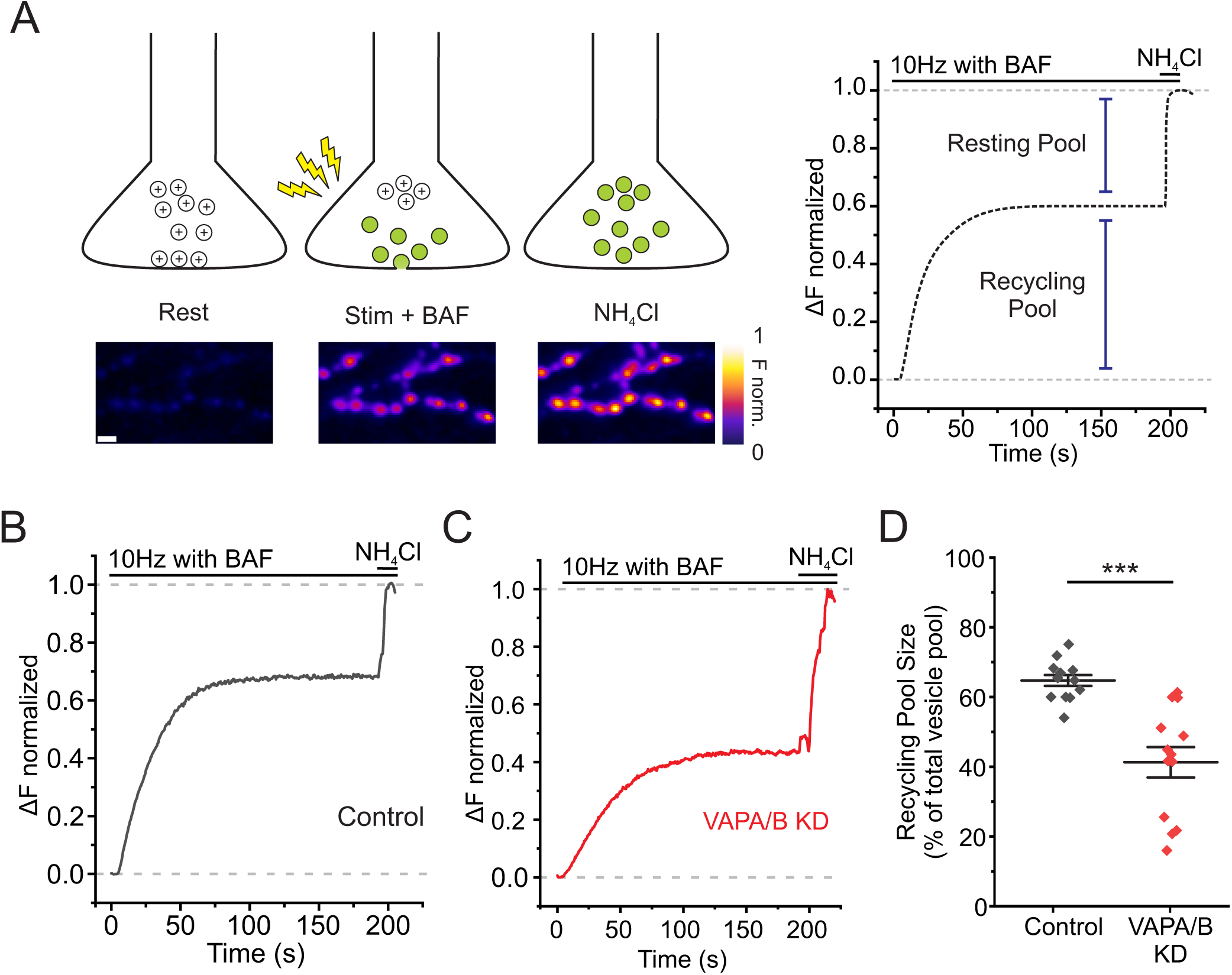
Loss of VAP shifts the distribution of vesicles from the recycling pool into the reserve pool. (A) Cartoon depicting experimental paradigm to determine recycling pool size using vG-pH, adapted from (83). During stimulation in the presence of Bafilomycin A1, vG-pH fluorescence represents cumulative exocytosis. Over the course of minutes during continuous 10Hz stimulation, all the vesicle in the bouton that are release-competent, comprising the RRP and recycling pool, will fuse (Left). Briefly washing NH_4_Cl reversibly collapses the proton gradient in vesicles and reveals the total vesicle pool and can be compared to the fraction of vesicles released during continuous stimulation to find the recycling and resting pool fractions (Right). (*B* and *C*) Representative fluorescent vG-pH traces for control (*B*) and VAPA/B KD neurons (*C*). (*D*) Mean values of recycling pool size in control and VAPA/B KD neurons (control, n = 10; VAPA/B KD, n = 12; \*\*\**P*<0.001 Student’s *t-*test). VAPA/B KD neurons have a smaller recycling pool size compared to control neurons, suggesting that mechanisms of vesicle mobilization are supported by VAP MCSs in the synapse.

## Discussion

Here we identify a presynaptic role for VAP as a molecular organizer of Ca²⁺ handling and synaptic vesicle availability during sustained activity. We show that VAP is required to sustain vesicle fusion during trains of stimulation (Fig. 2). Although loss of VAP does not measurably alter global cytosolic Ca²⁺ signals evoked by electrical activity, it markedly disrupts activity-dependent Ca²⁺ accumulation within the ER (Fig. 3) and mitochondria (Fig. 4). Notably, the vesicle-fusion and ER Ca²⁺ phenotypes closely phenocopy disruption of the plasma-membrane binding partner of VAP, Kv2 (Fig. 4E–F and Fig. S1), consistent with a functional role for Kv2/VAP ER–PM membrane contact sites (MCSs) in presynaptic boutons.

Mechanistically, we find that the dominant defect underlying reduced exocytosis is impaired mobilization of vesicles into the fusion-competent recycling pool: in VAPA/B knockdown neurons, a substantial fraction of vesicles remains sequestered in the reserve pool and are unavailable for release during prolonged stimulation (Fig. 6). These results align with prior work reporting reduced synaptic vesicle internalization after VAP knockdown and are consistent with genetic studies at Drosophila neuromuscular junctions (NMJs), where VAP perturbation is associated with altered presynaptic Ca²⁺ dynamics and impaired ER Ca²⁺ release (62). Together, our findings extend these observations by supporting a model in which VAP/Kv2-organized ER–PM MCSs shape presynaptic Ca²⁺ distribution and thereby regulate vesicle mobilization upstream of fusion at the active zone.

Ca^2+^ plays distinct yet equally critical roles in both the fusion and mobilization of synaptic vesicles (63–66). Vesicle fusion and neurotransmitter release are initiated by high Ca^2+^ concentrations within microdomains formed by Cavs in the active zone. These Ca^2+^ microdomains activate Ca^2+^ sensors to initiate vesicle fusion. However, elevated intracellular Ca^2+^ also accelerates the recruitment of release-ready vesicles (67, 68), thereby sustaining vesicle release during bursts of electrical activity that are critical for short-term plasticity. Importantly, Ca^2+^-dependent vesicle replenishment responds to much different levels of intracellular Ca^2+^ than classical vesicle fusion machinery, yet powerfully influences synaptic strength (69, 70). This effect is often attributed to Ca^2+^ diffusion from the active zone elevating global Ca^2+^. However, vesicle replenishment is selective and does not appear to depend on proximity to the active zone (71, 72). This finding strongly suggests that Ca^2+^ signaling within the vesicle pools is localized and specific for the source of Ca^2+^ rather than diffusion from the active zone alone. This raises the possibility of a secondary Ca^2+^ source organized by the ER within the terminal that may be critical for synaptic function.

The relationship between Kv2/VAP organization and vesicle recruitment is also consistent with findings in other secretory cell types. Kv2 has a non-conducting role in promoting exocytosis in neuroendocrine cells, including chromaffin cells (73) and pancreatic β-cells (74), where impaired dense-core vesicle recruitment is a prominent phenotype. Although this function was initially attributed to Kv2 C-terminal interactions with syntaxin, a truncated Kv2.1 construct that retains electrical function and syntaxin binding but cannot form clusters failed to enhance granule recruitment or exocytosis (74). These results support the idea that Kv2 clustering—and by extension VAP-dependent organization of Kv2-containing membrane domains—plays a broader role in coordinating vesicle recruitment beyond synaptic terminals.

Although VAP is frequently found at ER-mitochondria MCSs, the functional significance of this localization in axons remains unclear. VAP has been implicated in stabilizing mitochondria in distal dendrites, in a Kv2-independent manner, whereas this role was not evident in axons (36), suggesting compartment-specific functions for VAP-containing MCSs (36). In our experiments, loss of VAP strongly reduced activity-evoked mitochondrial Ca²⁺ uptake, yet did not produce a clear impairment in ATP-dependent processes such as endocytosis (Fig. S1), consistent with the possibility that residual mitochondrial Ca²⁺ entry is sufficient to support energy production under these conditions. Axonal mitochondria can acquire Ca²⁺ directly from the cytosol through the MCU complex and MICU regulators, including MICU3, which enhances activity-dependent uptake in neurons (50). In agreement with this, blocking ER Ca²⁺ uptake via SERCA inhibition did not alter mitochondrial Ca²⁺ signals in control cells (Fig. 5G–H). Nonetheless, disrupting Kv2/VAP MCSs markedly reduced mitochondrial Ca²⁺ uptake efficiency, suggesting that mitochondrial Ca²⁺ entry may exhibit source preference within the bouton—potentially via spatial organization of Ca²⁺ flux near organelle interfaces. Reduced mitochondrial Ca²⁺ sequestration could, over time, compromise synaptic health by diminishing presynaptic Ca²⁺ buffering capacity (75–77). Alternatively, reduced mitochondrial Ca²⁺ uptake may partly reflect decreased energetic demand if fewer vesicles participate in release, given that the vesicle cycle is a major presynaptic ATP sink (56).

This ability of ER MCSs to serve as a scaffold for Ca^2+^ signaling is well established in other excitable cells, such as in muscle, where Cavs (Cav1 isoforms), RyRs, and junctophilin enable muscle contraction (78, 79). In the neuronal soma, Kv2/VAP MCSs are enriched in L-type Cavs through a physical interaction within the Kv2.1 C terminus (46) that influences the probability of L-type Cav opening (14) and mediates excitation-transcription coupling (46). This ability for ER MCSs to coordinate Ca^2+^ signaling is not specific to Cav1s, but extends to Cav2s in heterologous cells (80) and in neurons, where junctophilin-containing MCSs are critical for CaMKII activation in dendrites (6). Our data suggest that ER/PM MCSs coordinate organelle Ca^2+^ signaling during electrical activity, and supports our hypothesis that this organization is present in presynaptic boutons. However, characterizing presynaptic MCSs present technical challenges. The diameter of presynaptic MCSs is <30nm, which often precludes immunostaining due to epitope accessibility, steric hindrance, and insufficient molecular density for fluorescence detection in fine structures like axons. Furthermore, MCSs exhibit molecular diversity (81), and it remains unclear which proteins beyond Kv2 and VAP occupy these junctions or whether distinct MCS subtypes exist presynaptically. Recent work demonstrates that STIM1 and ORAI proteins occupy different MCSs than Kv2 in neuronal somas (82), suggesting specialization among MCS subtypes that may also extend to presynaptic terminals.

Finally, while loss of VAP is implicated in rare familial forms of amyotrophic lateral sclerosis (ALS), the extent to which our findings map onto motor neuron pathology remains to be established. ALS is a progressive and fatal neurodegenerative disease characterized by degeneration of upper and lower motor neurons, leading to muscle denervation, atrophy, and respiratory failure. Although our cultured rat hippocampal neuron model does not recapitulate motor neuron-specific disease processes, it provides a robust system for defining fundamental presynaptic mechanisms. Notably, the synaptic transmission and vesicle mobilization impairments observed with VAP depletion (Figs. 2 and 6) parallel features reported in motor neuron disease models, supporting the possibility that disrupted ER–PM organization contributes to synaptic vulnerability in disease contexts. Overall, our work provides cellular-level evidence that VAP and Kv2 organize ER–PM MCSs that support synaptic transmission by coordinating presynaptic organelle Ca²⁺ handling and vesicle mobilization.

## Methods

### Cell Culture and Transfection

Primary neurons from Sprague-Dawley rats of either sex on postnatal day 0 or 1 were cultured for all experiments. Hippocampal CA1 to CA3 regions with the dentate gyri removed were harvested, tissue was dissociated into single cells with bovine pancreas trypsin, and cells were plated onto poly-L-ornithine-coated coverslips inside a 6-mm cloning cylinder as previously described (84). Ca^2+^ phosphate-mediated DNA transfection was performed on cultures at 5 to 6 DIV. All experiments were performed on mature neurons between 14 and 24 DIV. To ensure reproducibility, experiments were performed on neurons from a minimum of three separate cultures. All protocols used were approved by the Institutional Animal Care and Use Committee at Dartmouth College and conform to NIH Guidelines for the Care and Use of Animals.

### Genetic Tools

The following constructs were used: mGreenLantern (Addgene #161912, hSyn promoter) (85), vGlut-pHluorin (43), jGCaMP8f (Addgene # 162376) (45), ER-GCaMP6-150 (Addgene #86918) (7), mito^4x^-GCaMP6f (Addgene # 127870) (50), mCherry-hVAPA (Higgs Lab, Dartmouth College), mCherry-hVAPB (Addgene # 108126) (86), mitoGFP (Higgs Lab, Dartmouth College), mRuby-Synapsin (de Juan-Sanz Lab, ICM-Paris Brain Institute). To reduce endogenous VAPA/B expression for live cell imaging, a tandem sgRNA plasmid (Addgene # 64073) (87) was designed against the following mRNA sequences, VAP-A sgRNA: AGACAGTCACGATTGACCCT; VAP-B sgRNA: CTGCGTGCGGCCCAACAGTG. To reduce endogenous Kv2.1 expression for live cell imaging, an shRNA plasmid was obtained from Origene (TR709776, variant A) (26). PTPIP51 shRNA was obtained from Origene (TR702178, variant A).

### Reagents

All chemicals were obtained from Sigma except for cyclopiazonic acid (CPA) (Alomone), bafilomycin-A1 (Tocris), and dantrolene sodium salt (Tocris).

### To validate the effectiveness of our KD strategy

Neurons were fixed with 4% paraformaldehyde and 4% sucrose in phosphate-buffered saline (PBS) for 10 min, permeabilized with 0.2% Triton X-100, and blocked with 5% goat serum in PBS for 30 min at room temperature. Neurons were then incubated with the VAPA/B antibody N479/107 (Neuromab) and GFP primary antibody (A10262, Invitrogen) overnight at 4°C. Cells were washed three times with PBS and incubated for 1 h with Alexa Fluor-conjugated secondary antibodies (Invitrogen).

### Widefield Live Cell Imaging

All experiments were performed at 35°C using a custom-built objective heater. Culture cells were mounted in a rapid-switching laminar flow perfusion and stimulation chamber on the stage of a custom-built epifluorescence laser microscope. The total volume of the imaging chamber was 100µl and neurons were perfused at a rate of 400µl/min in a modified Tyrode’s solution containing the following (in mM): 119 NaCl, 2.5 KCl, 2 CaCl_2_, 2 MgCl_2_, 25 HEPES, and 30 D-glucose with 10µM cyanquixaline (Sigma-Aldrich) and 50 µM APV (Sigma-Aldrich). Images were obtained using a Zeiss Observer Z1 equipped with an EC Plan-Neofluar 40X 1.3 numerical aperture (NA) oil immersion objective. Images were captured on IXON Ultra 897 Electron Multiplying Charge Coupled Device (EMCCD) (Andor) that was cooled to −80°C by an external liquid cooling system (EXOS). All excitation light occurred via OBIS lasers (Coherent). Action potentials (APs) were evoked by passing 1-ms current pulses yielding fields of ∼12 V/cm^2^ via platinum/iridium electrodes. Timing of stimulation was delivered by counting frame numbers from a direct readout of the EMCCD rather than time itself for more exact synchronization using a TTL from nStrobe (Prizmatix).

### Confocal Live Cell Imaging

Live cells were imaged on a Nikon Ti2-inverted CSU-W1 SoRa spinning disk confocal microscope using a 40X 1.3 NA oil immersion objective, excitation using 488 nm (mGreenLantern-Kv2.1, Fig 1; mitoGFP, Fig 5) and 561nm (mRuby-Synapsin, Fig 5) wavelength and imaged with 100-400ms exposure time. Z-stack with 0.3mm steps were taken of an individual cell. Z-stack was projected using ImageJ (NIH).

### GCaMP Ca^2+^ measurements

Cells were illuminated by an OBIS 488nm laser from 2-10mW (Coherent) with ET470/40x, ET525/50 m, and T495Ipxr filters (Chroma). We repeated and averaged three trials to measure AP train stimulation-induced responses. jGCaMP8f and mito^4x^-GCaMP6f fluorescence was recorded with a 9.8ms exposure time and images were acquired at 100 Hz. ER-GCaMP6-150 fluorescence was recorded with a 19.8ms exposure time and images were acquired at 50 Hz. GCaMP peak fluorescence (ΔF/F_0_) for each stimulation was found by averaging the 5 highest points divided by the average pre-stimulus fluorescence time points.

### Vesicle fusion measurements

vGlut-pHluorin (vG-pH) fluorescence was recorded with a 9.8ms exposure time and images were acquired at 100 Hz. Cells were illuminated by an OBIS 488-nm laser at 6 to 10 mW (Coherent) with ET470/40x, ET525/50 m, and T495Ipxr filters (Chroma). Cells were bathed in 50mM NH_4_Cl to neutralize vesicle pH at the end of each experiment to quantify vesicle fusion. Peak exocytosis for each stimulation was found by averaging 5 frames (5 ms) after train stimulation. To quantify endocytosis, vG-pH was recorded with 500ms exposure time and images were acquired at 2Hz. A single exponential decay was fit to the corresponding fluorescence 2s after stimulation ended to account for surface dwell time to capture pure endocytosis (57). Endocytosis measurements included in the present dataset reached at least an R-squared of 0.95 for exponential decay.

### Image and Data Analysis

Images were analyzed in ImageJ using a custom-written plugin (imagej.net/ij/plugins/time-series.html). To quantify fluorescence, we selected 1.4-µm-diameter circular regions of interest (ROIs) from ΔF images of each experiment, recentering on the brightest pixel within the ROI in the ΔF image. A minimum of 30 ROIs were averaged together for a single cell. ROIs were selected based on differential fluorescence images (F_peak_ – F_baseline_) of vesicle fusion, Ca^2+^, rather than morphology, to define a presynaptic terminal. All statistical data are presented as means ± SEM (n = number of neurons) and all experiments were performed on more than three independent cultures. Quantification of vesicle fusion was obtained by normalizing the fluorescence change in response to stimulation to the total number of vesicles measured by application of 50mM NH_4_Cl. Bafilomycin A1 and NH_4_Cl were analyzed by mean value of plateau regions. Images were pseudocolored, cropped, and adjusted for contrast and brightness. Details specific to each experiment are presented in results or figure legends.

### mGreenLantern-Kv2.1 clustering analysis

To quantify the degree of Kv2.1 clustering, we drew a line scan with a width of 5 pixels (0.16µm/pixel x 5 = 0.8µm wide) around the soma at a single z plane, through the soma’s midpoint. We then calculated the background-subtracted average fluorescence and determined its coefficient of variation (CV).

### Mitochondria-Synapse quantification

To quantify mitochondria and synapse distribution along axons, we drew line scans with a minimum length of 50µm and a width of 10 pixels (1.6µm; 0.16µm/pixel). We calculated the Spearman correlation coefficient between the corresponding fluorescence intensities of the two channels. To identify synaptic boutons (mRuby-Synapsin) and mitochondria (mitoGFP), we applied intensity thresholds of 20x and 100x the basline standard deviation, respectively. We then quantified peak characteristics (FWHM) using Peak Analyzer in OriginPro with a local maximum setting of 4 pixels (∼500nm) and the thresholds defined above. To determine the percentage of synapses with proximal mitochondria, we took a 1µm radius around the center of each synapse and scored whether a mitochondrion based on the threshold criteria was present within this region.

### Quantification and Statistical Analysis

Error bars indicate mean ± SEM unless otherwise noticed. Statistical analyses were performed in Excel and Origin Pro. We used the paired two-sample for means *t-*test for paired results. Normally distributed data were processed with the Student’s *t-*test for two independent distributions. We used the One-way ANOVA for results with more than two experimental groups and *post hoc* Tukey’s test for pairwise comparisons.

### Data Availability

All study data are included in the article and/or *S1 Appendix*

## Acknowledgments

This work was supported by NIH National Institute of Neurological Disorders and Stroke Grants F31NS141318 (C.D.P) and 1RO1NS112365-01A1 (M.B.H), the National Science Foundation Career IOS 1750199, NIH award S10OD032310 to Dr. Yashi Ahmed and Bing He for purchase of the super-resolution spinning disk confocal, Life Sciences Light Microscopy Facility at Dartmouth College, bioMT through NIH grant P20-GM113132, DartCF through NIH grant P30-DK117469, and NCI Cancer Center Support Grant P30 CA023108. We thank Dr. Michael Tamkun (Colorado State University) and Dr. Henry Higgs (Dartmouth College) for plasmids used in these experiments, and Dr. Andrew Coleman for designing our VAP sgRNA constructs. We also thank members of the Hoppa lab including Emily Dueñas Cornell du Houx, In Ha Cho, Aman Aberra, Lauren Panzera, Kiera Schwarz, Matthew Bazan, and Michelle Gleason for their careful reading of the manuscript. Finally, Dr. Michael Tamkun provided careful reading and valuable feedback of an early version of the manuscript.

---

**Figure S1.**
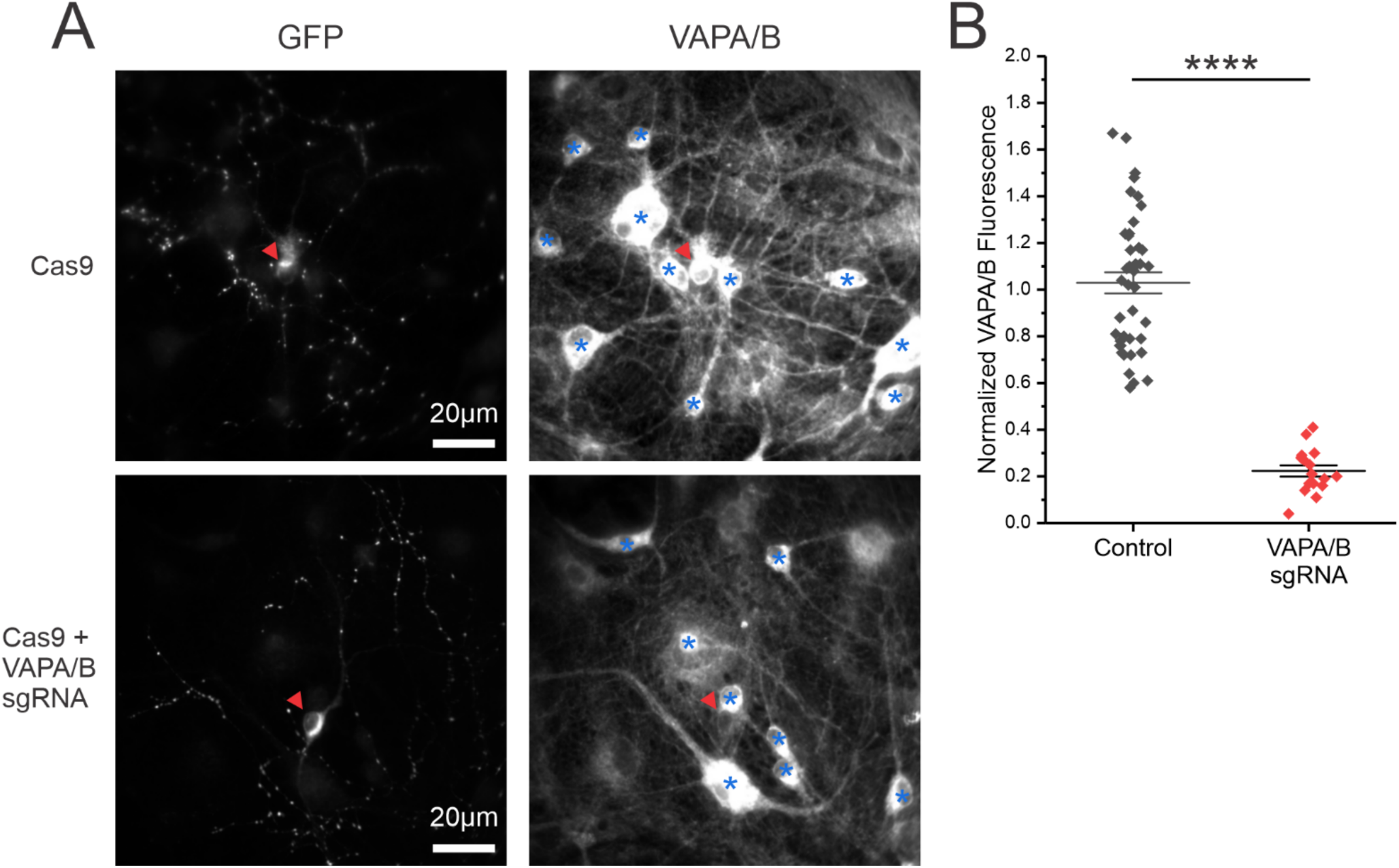
VAPA/B expression is reduced by sgRNA KD. VAPA/B sgRNA was designed to efficiently deplete both isoforms of VAPA/B to test roles in synaptic transmission. Plasmid encoding sgRNA was delivered using Ca^2+^ phosphate-based transfection to a small fraction of neurons (<1% of total neurons). This allows for comparison of protein expression levels of each cell with adjacent untransfected neurons to determine transfection efficiency. (**A**) Cultured hippocampal neurons were immunostained for GFP and endogenous VAPA/B. Red arrowheads marks a neuron co-transfected with GFP and VAPA/B sgRNA that is depleted of endogenous VAPA/B. Blue asterisks indicated untransfected controls. By taking the fluorescence intensity of transfected cells (red arrowheads) and dividing the untransfected cells (blue asterisks), we can quantify relative VAPA/B expression levels. Minimum of 4 control neurons in each field of view. (**B**) Quantification of immunostained fluorescence normalized to control (control neurons, n = 42; VAPA/B KD neurons, n = 16; *P* < 0.0001, Student’s *t*-test).

**Figure S2.**
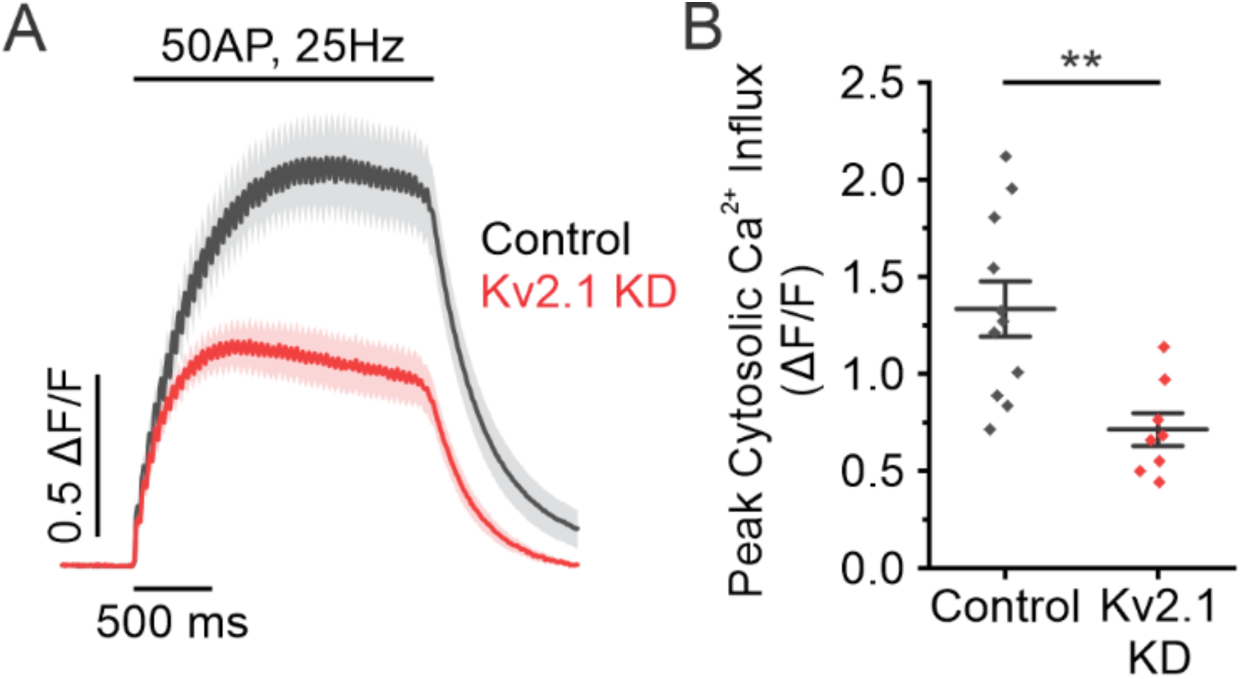
Loss of Kv2.1 impairs cytosolic Ca^2+^ influx. (A) Average fluorescent traces of jGCaMP8f for control neurons and Kv2.1 KD neurons in response to 50 AP stimulation delivered at 25 Hz. (B) Quantification of peak fluorescence for control and Kv2.1 KD neurons stimulated with 50 AP delivered at 25Hz (control neurons, n = 11; Kv2.1 KD neurons, n = 8; *P* < 0.01, Student’s *t-*test).

**Figure S3.**
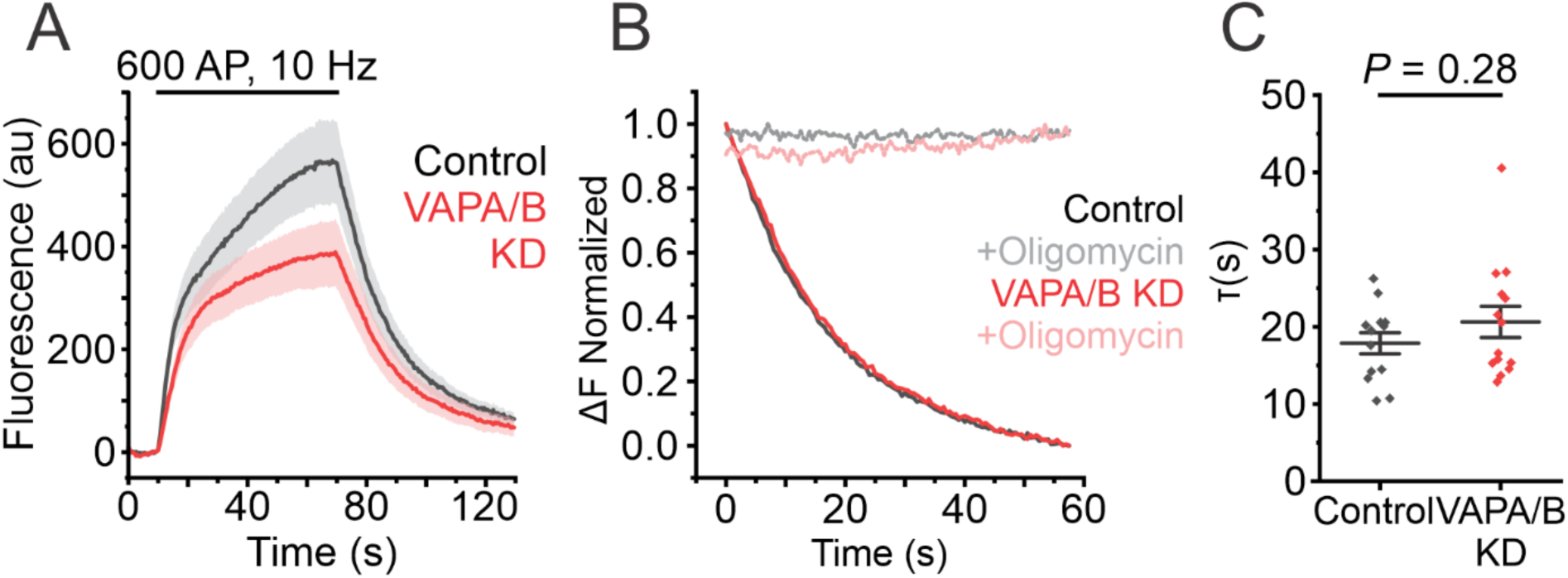
VAPA/B KD neurons do not show any defects in endocytosis as a proxy for ATP availability. (A) Average fluorescent traces of vGlut-pHluorin in control cells and VAPA/B KD cells in response to 600AP delivered at 10Hz in Tyrode’s imaging solution containing 1.25mM lactate and pyruvate. (B) Average normalized fluorescent traces comparing exponential decay for modified Tyrode’s imaging solution containing Oligomycin A to show that there is no residual glucose present when switching between Tyrode’s solutions. Exponential decay are fit 2s after the end of stimulation to account for vGlut-pHluorin surface dwell time (57). (*C*) Endocytosis time constants (tau, 1) of vGlut-pHluorin control neurons and VAPA/B KD neurons following 600 AP, 10 Hz stimulation in Tyrode’s solution containing lactate and pyruvate (control neurons, n = 13; VAPA/B KD, n = 14; *P* = 0.28, Student’s *t-*test).

